# Mesoscale molecular assembly is favored by the active, crowded cytoplasm

**DOI:** 10.1101/2023.09.19.558334

**Authors:** Tong Shu, Gaurav Mitra, Jonathan Alberts, Matheus P. Viana, Emmanuel D. Levy, Glen M. Hocky, Liam J. Holt

## Abstract

The mesoscale organization of molecules into membraneless biomolecular condensates is emerging as a key mechanism of rapid spatiotemporal control in cells^1^. Principles of biomolecular condensation have been revealed through *in vitro* reconstitution^2^. However, intracellular environments are much more complex than test-tube environments: They are viscoelastic, highly crowded at the mesoscale, and are far from thermodynamic equilibrium due to the constant action of energy-consuming processes^3^. We developed synDrops, a synthetic phase separation system, to study how the cellular environment affects condensate formation. Three key features enable physical analysis: synDrops are inducible, bioorthogonal, and have well-defined geometry. This design allows kinetic analysis of synDrop assembly and facilitates computational simulation of the process. We compared experiments and simulations to determine that macromolecular crowding promotes condensate nucleation but inhibits droplet growth through coalescence. ATP-dependent cellular activities help overcome the frustration of growth. In particular, actomyosin dynamics potentiate droplet growth by reducing confinement and elasticity in the mammalian cytoplasm, thereby enabling synDrop coarsening. Our results demonstrate that mesoscale molecular assembly is favored by the combined effects of crowding and active matter in the cytoplasm. These results move toward a better predictive understanding of condensate formation *in vivo*.

---

Cells are highly crowded, with up to 40% of cellular volume excluded by macromolecules^4,5^. This high excluded volume can inhibit molecular motion, but on the other hand can entropically favor assembly through depletion attraction forces^6,7^. The majority of cytoplasmic volume is taken up by mesoscale (10 – 1000 nanometer diameter) particles^5^. This means that the effects of crowding strongly affect the behavior of mesoscale particles and assemblies, while having less impact on nanoscale processes because nanoscale particles can move relatively freely between mesoscale crowders, but mesoscale particles cannot. Studies have shown that macromolecular crowding can change biochemical reaction kinetics, protein conformations, and motor functions^7–9^.

The cell also contains elastic networks that constrain and organize the cell interior. These include the actomyosin cytoskeleton in the cytoplasm^10^ and chromatin in the nucleus^11^. The presence of these networks and the high concentration of particles together make the intracellular environment viscoelastic. This contrasts with simple buffer solutions, which are only viscous.

Finally, cells are non-equilibrium open systems, and use adenosine triphosphate (ATP)-dependent cellular activities to maintain a non-equilibrium steady state by exchanging energy, information and material with the extracellular environment, thereby locally reducing entropy^12^. Overall, the intracellular environment is highly complex, and its impact on the assembly of membraneless biomolecular condensates remains largely unexplored.

The assembly of membraneless biomolecular condensates bridges length scales between the nanoscale and mesoscale, where nanometer diameter molecules come together to form higher-order structures of tens to thousands of nanometers in diameter^13^. This wide range of length- and time-scales makes it difficult to predict how the crowded, active cellular environment will affect biomolecular condensate formation. Several studies have focused on the impact of elastic mechanical properties on condensate growth^14–17^. For example, elastic chromatin mechanics has been shown to frustrate the growth of nuclear condensates^16,17^. However, the combined impacts of macromolecular crowding, elastic networks, and non-equilibrium cellular activities on condensate formation are less well understood.

It is difficult to derive general physical principles from the study of endogenous condensates because these systems are formed through complex coacervation of many molecules. Furthermore, these components are often dynamically altered by post-translational regulation, the details of which are typically unknown. Thus, when perturbing intracellular environments, it is difficult to fully attribute structural changes in endogenous condensates to only biophysical cues, since biological functional changes associated with perturbations can also lead to structural changes in endogenous condensates. To overcome these issues, we developed an orthogonal synthetic intracellular condensate system called synDrops. synDrops adapted a previous approach to create a molecular condensate of well-defined geometry^18^, but adds the ability to chemically induce the interaction of components.

We successfully induced synDrop formation in both budding yeast *S. cerevisiae* cells and mammalian cervical cancer HeLa cells. Complementary to the experimental system, we also developed two independent agent-based molecular dynamics models to simulate synDrops within cellular environments from first principles. Combining experiments and simulations, we show that macromolecular crowding facilitates the nucleation process while inhibiting the growth phase of condensate dynamics. However, ATP-dependent cellular activity promotes growth by assisting long-range structural rearrangements. In conclusion, we found that the assembly of mesoscale biomolecular condensates is favored by the crowded and active cellular environment.

## Droplet dynamics can be captured using inducible synDrops in cells and *in-silico*

SynDrops are composed of two protein components, each of which has three modular domains. We based our design on the Flory-Stockmeyer theory^19^, which governs polymer network growth. Multivalency is essential for the formation of mesoscale condensates through phase separation^20–24^. We used homomultimerizing domains to create multivalency in our system (Fig 1a). One component uses a homohexamer multimerization domain (PDB: 3BEY), and the other uses a homodimer domain (PDB: 4LTB). The two components interact in *trans* through two halves of an inducible heterodimeric binding interaction, enabling kinetic analysis. Importantly, the dimerization domain is a 19 nm long, stiff, antiparallel coiled-coil. Since the distance between interaction surfaces on the hexamer is approximately 6 nm, the dimer is sterically prevented from interacting with the same hexamer component more than once. Thus, geometric constraints strongly favor the expansion of synDrop molecular networks, which greatly simplifies simulation and physical analysis compared to other synthetic systems^22–24^ (Fig 1a).

**Fig. 1.**
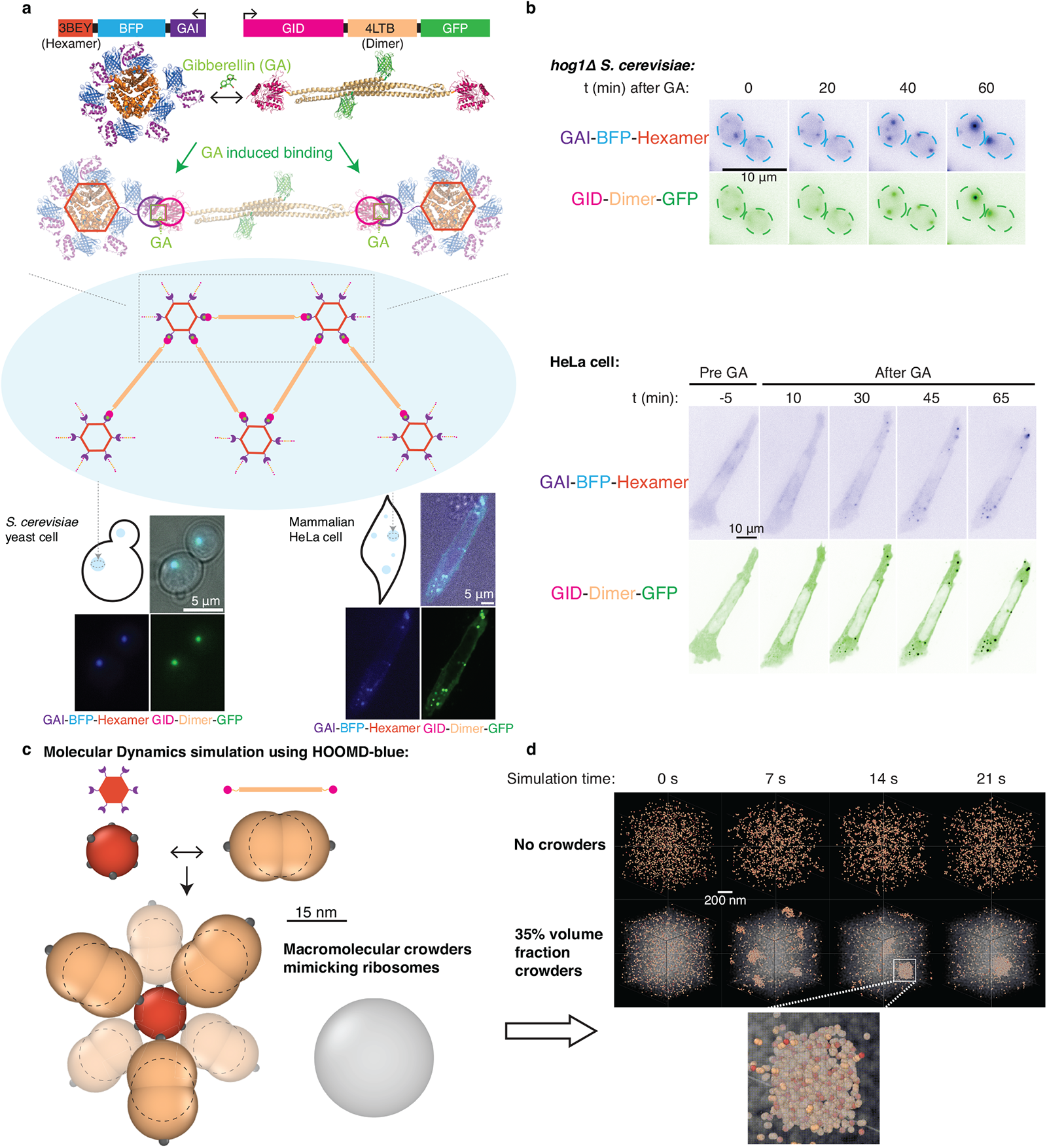
synDrops enable the analysis of condensate formation kinetics in *S. cerevisiae* and mammalian cells, and are amenable to simulation. **a**, synDrops are inducible synthetic condensates composed of two proteins. Top) Gene and crystal structures of the two components. Each protein has three domains: an oligomerization domain (3BEY: Hexamer or 4LTB: Dimer), an inducible interaction domain (GAI or GID) and a fluorescent protein (BFP or GFP). Gibberellin (GA) induces binding between GAI and GID favoring the formation of a mesoscale molecular network (middle cartoon), as shown in representative images of both *S. cerevisiae* yeast cells and mammalian HeLa cells (bottom). **b,** GA addition leads to synDrop formation. Time course of synDrop formation in *hog1*Δ *S. cerevisiae* yeast cells, and mammalian HeLa cells. Scale bar, 10 *μm* **c,** Schematic of molecular dynamics model for synDrop assembly. The hexamer component is represented by a sphere with six uniformly-distributed binding sites; the dimer component is represented as a rod-like structure formed from three overlapping spheres with two binding sites positioned on opposite ends. A third component with no binding sites mimics ribosomes as macromolecular crowders. **d,** Representative images from HOOMD-blue MD simulations of synDrops system over time without crowders (top) and with 35% volume fraction of crowders (bottom).

The inducible binding domains are the plant GAI (Gibberellin insensitive DELLA proteins) and GID (Gibberellin Insensitive Dwarf 1) domains. These domains undergo a heterotypic interaction that is potentiated in the presence of the plant hormone Gibberellin (GA)^25^ (Fig. 1a). GAI is truncated to a minimum dimerization domain^26^. Adding GA increases the affinity between the two synDrop components and triggers synDrop formation.

We co-expressed these two proteins, in yeast cells by integrating two plasmids into the genome or in mammalian HeLa cells through transient transfection. Since the ratio between these two proteins is essential for condensate formation^18^, we adapted our plasmid design for expression in mammalian cells by combining the two genes onto the same plasmid and separating them by a P2A self-cleavage sequence (Extended data Fig. 1a). P2A self-cleavage peptide can trigger ribosome skipping forming a glycyl-prolyl peptide bond at the C-terminus of P2A sequence during translation, resulting in two independent proteins^27^ at equal expression levels. After the proteins were expressed within cells, we added GA into the cell media and observed synDrop dynamics at different time points after GA addition (Fig. 1b). Similar to the previous reports^18^, synDrops were spherical and were observed to fuse suggesting they had liquid-like material properties (Extended data Fig. 1b).

Our *in vivo* system enables detailed analysis of mesoscale assembly, but cannot easily report on the microscopic protein interactions that underpin this process. Therefore, we developed two independent agent-based molecular dynamics (MD) platforms to provide complementary information *in silico*. The first simulation setup used a HOOMD-blue engine^28,29^ combined with a dynamic bonding plugin that we previously developed^30^ (Fig. 1c and 1d), and the second used a custom-developed Java program (Extended data Fig. 1c and 1d). In MD simulations with HOOMD-blue, we modeled the hexamer as a single sphere with six uniformly-distributed binding sites, and the dimer as a rod-like structure formed from three spheres with two binding sites positioned on opposing sites of the two outer spheres (Fig. 1c). In the Java MD simulations, we modeled the hexamer and dimer as spheres with 6 or 2 binding sites respectively (Extended data Fig. 1c). The sizes of these simulated structures were chosen based on crystal structure information. In addition, we included a third agent to mimic ribosomes, which are the dominant macromolecular crowders in the cytoplasm. This agent was a 30 nm diameter sphere with no binding interactions. The formation of synDrops was simulated with or without crowders under equilibrium conditions (Fig. 1d and Extended data Fig. 1d). There is a discrepancy in the time scales of synDrops formation in experiments and simulations. This may be for several reasons, including that the cell is around 200 times larger than our simulated system. In summary, we have developed the synDrop system both in cells and *in silico*, allowing us to address how the intracellular environment affects the assembly of mesoscale condensates.

## Macromolecular crowding promotes nucleation but inhibits droplet growth

We examined synDrop kinetics within cells under various conditions using fluorescence microscopy. We first characterized GA induced droplet dynamics in yeast cells by quantifying the average number of droplets per cell, as well as droplet total intensity (see methods). Changes in droplet total intensity are indicative of changes in cluster size and/or its molecular density. Droplet growth occurred in two phases. First, there was a nucleation phase, during which the average droplet number per cell increased. Subsequently, droplets grew by fusion and coarsening, leading to a decrease in droplet number and an increase in droplet intensity.

Under control conditions, we observed an increase in droplet number within the first 10 minutes after inducing binding interactions by the addition of GA (gray curve, Fig. 2a). However, the total intensities of those newly formed droplets did not grow in this period (Fig. 2b). This suggests that droplets initially nucleate locally but do not grow substantially. After 10 minutes, the droplet number started to decrease and the intensity started to increase. This corresponds to a growth phase where droplet sizes become larger, mainly through droplet coalescence. Thus, the synDrops system forms droplets by nucleation and growth, which has been suggested as the most common mechanism of endogenous condensate formation^31,32^.

**Fig. 2.**
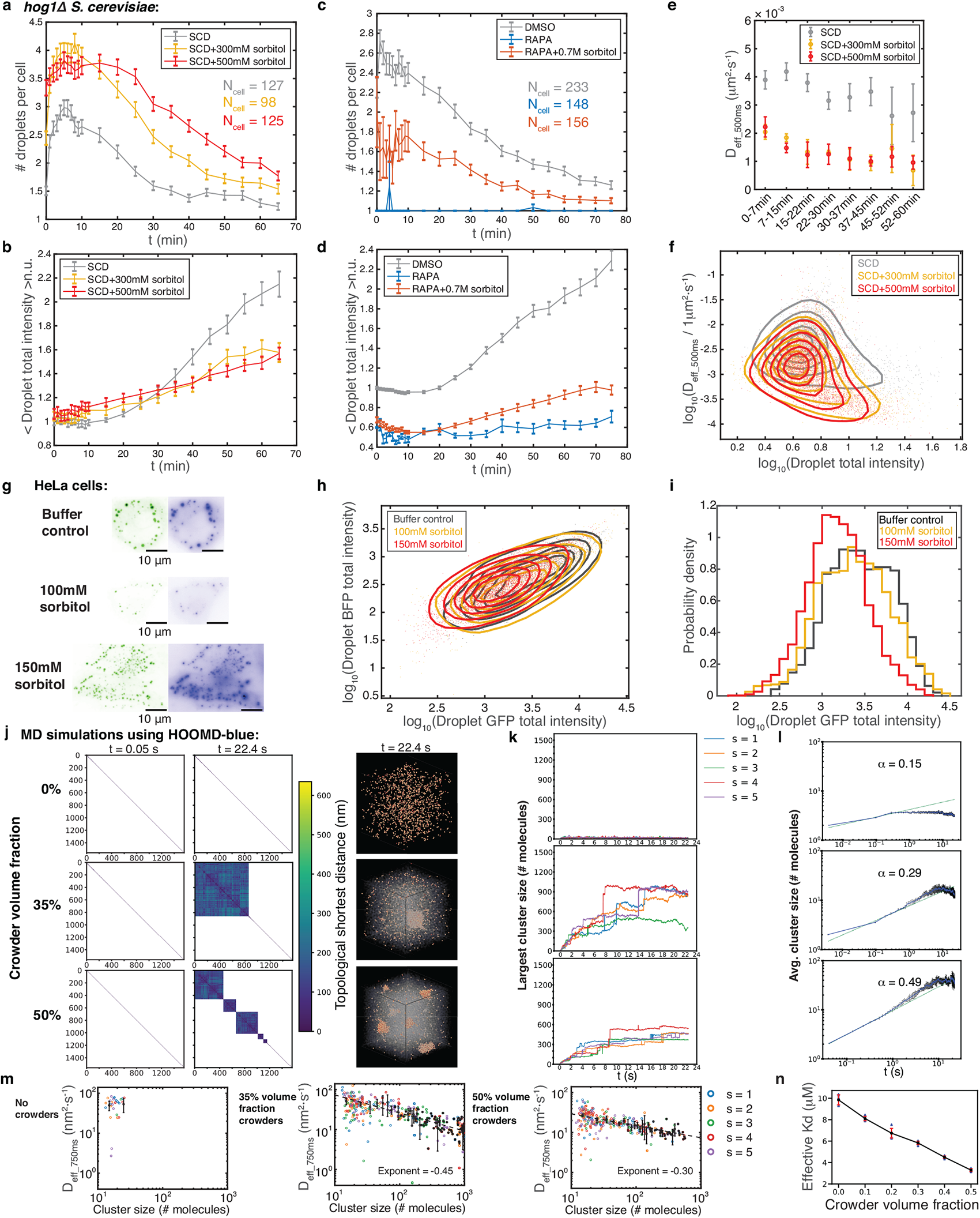
Increasing molecular crowding promotes synDrop nucleation but inhibits growth. **a**, Averaged number of droplets per cell (*hog1*Δ *S. cerevisiae*) for one hour after synDrop induction with GA for control and two osmotic compression conditions. Error bars are standard error of mean (SEM). **b,** Total intensities of droplets normalized to the averaged value of control cells at 0 min. Data are mean ± SEM. **c,** Averaged number of droplets per cell and **d,** normalized droplet total intensity over one hour comparing control (DMSO) and decreased molecular crowding (Rapamycin, RAPA) conditions, as well as RAPA treatment with recovered molecular crowding (RAPA + 700 mM sorbitol). Data are mean ± SEM **e,** Median diffusivity of droplets at different time points post-induction in the same conditions as **a** & **b**. Error bars are SEM. **f,** Density plot of droplet diffusivity versus total intensity at all time points post-induction. **(g-i),** synDrops formed in mammalian HeLa cells after 1-2 hours of induction comparing control to osmotic compression conditions (100 mM or 150 mM sorbitol). **g,** Representative images, scale bar 10 *μm*. **h,** Phase diagram of synDrop formation as a function of BFP and GFP total intensities. **i,** Histogram of droplet GFP total intensities. **(j-n),** Analyses of MD simulations using HOOMD-blue comparing conditions with 0%, 35% and 50% volume fractions of crowder: **j,** Graph theory analyses (left) of cluster formation at early and later times with corresponding simulation renderings (right). **k,** Number of molecules within the largest cluster over time from five replicate simulations, denoted by different colors. **l,** Averaged cluster size (number of molecules) over time from five replicate simulations. The dashed line represents the power law fit for the initial 0.5 s, exponent denoted as *α*. Error bars are standard deviation (SD) of the averaged values of the five repeats. **m,** Cluster diffusivity versus cluster size (number of molecules) on the log-log scale. The black data points represent the mean of averaged values from five repeats, and the error bars correspond to the SD among these averaged values. The dashed black line represents the linear fit on the log-log scale and the fitted slope is labeled as the exponent. **n,** Effective dissociation constants (Kd) of a simplified monovalent system as a function of crowder volume fraction. Error bars are SD from five repeats.

We next explored the effects of macromolecular crowding on synDrop formation. Osmotic compression of cells can increase macromolecular crowding. However, wild-type yeast can rapidly balance external osmotic pressure by producing glycerol^33^. To circumvent the osmo-adaptation in yeast cells, we used *hog1*Δ yeast cells (Fig. 1b). Hog1p is a key regulator kinase, required for rapid glycerol accumulation in yeast cells^33^. Therefore, deletion of the *HOG1* gene prevents rapid osmo-adaptation allowing us to more precisely tune molecular crowding; we used *hog1*Δ *S. cerevisiae* cells in this study unless otherwise stated.

After osmotic compression, the initial number of droplets nucleated was increased compared to control (yellow and red curve, Fig. 2a). However, the subsequent increase in droplet intensities was suppressed (yellow and red curve, Fig. 2b). Similar results were also obtained in WT yeast cells before osmo-adaptation started to have an effect (Extended data Fig. 2a). This suggests that macromolecular crowding promotes synDrop nucleation but inhibits growth.

Besides increasing macromolecular crowding, osmotic compression also increases the concentrations of the protein components (Extended data Fig. 2b). We took advantage of the noise in protein expression levels in single cells to evaluate the relative importance of increased protein concentration versus increased molecular crowding in changing synDrop dynamics. We first grouped cells in each condition (control and osmotic compression conditions) into four quantiles based on their cellular mean pixel intensities before GA induction, indicating protein expression levels. By quantifying the average number of droplets per cell within different cell quantiles for each condition, we observed that, in all conditions, the number of droplets per cell was higher in cell groups with higher intensities (4th quartile) and lower in cell groups with lower intensities (1st quartile) when compared to the overall average for all cells (Extended data Fig. 2c). This confirms that protein concentrations affect synDrop formation. However, if we selected control and compressed cells with the same range of protein concentrations, we obtained qualitatively similar results to Fig. 2a and Fig. 2b (Extended data Fig. 2d). These results indicate that the effects of osmotic compression on synDrop assembly kinetics is mainly due to increased macromolecular crowding.

We next sought additional means to change macromolecular crowding. It has been shown that ribosome concentrations are tuned through the TORC1 (target of rapamycin complex 1) pathway^5^. Inhibition of TORC1 using rapamycin reduces ribosome biogenesis and increases ribosome degradation, leading to lower ribosome concentration and therefore reduced macromolecular crowding. We treated yeast cells with either rapamycin or DMSO (solvent control) for 2 hours. We found rapamycin-treated cells did not form droplets after GA induction (Fig. 2c, 2d and Extended data Fig. 2e). However, GFP signal was reduced after rapamycin treatment (Extended data Fig. 2g). Thus, rapamycin treatment reduced the concentration of the dimer component, likely due to increased cell size and decreased protein translation upon TORC1 inhibition^34^. We again wished to determine how changes in macromolecular crowding and in protein concentration each impacted synDrop assembly. To achieve this, we sought to restore normal macromolecular crowding in rapamycin-treated cells. We leveraged a microrheology approach with Genetically Encoded Multimeric nanoparticles (GEMs) to quantify crowding^5^. We found that macromolecular crowding was decreased in rapamycin-treated cells, consistent with previous reports^5^ (Extended data Fig. 2f). We then osmotically compressed (Extended data Fig. 2f), and found that 0.7M sorbitol restored macromolecular crowding of rapamycin-treated cells to the level of control cells (Extended data Fig. 2f). However, this level of osmotic compression barely altered protein concentrations (Extended data Fig. 2g). In these conditions, we found that synDrop formation was recovered (Fig. 2c, 2d and Extended data Fig. 2e), but was somewhat less robust compared to control. These results further indicate that macromolecular crowding is crucial for synDrop assembly and protein concentrations are also important.

We next examined the mechanisms underlying the inhibition of droplet growth by macromolecular crowding. Droplets can grow in two ways: the first is through droplet coalescence^35^ and the other is through Ostwald ripening^36^. However, droplet coalescence has been suggested as the dominant mechanism for droplet growth in biological systems^16^. In this mechanism, the rate of droplet growth depends on the collision rate between two smaller droplets, which in turn depends on the diffusivities of these droplets^16,35^. We therefore hypothesized that macromolecular crowding inhibits droplet growth by reducing droplet diffusivities. To test this hypothesis, we quantified synDrop diffusivities through particle tracking in both control conditions and after increasing macromolecular crowding. Since droplet size increases with time, we analyzed droplet diffusivities at different time points after induction. Similar to previous results (Fig. 2b), droplet intensity profiles showed the same trends: droplet intensities were lower in conditions of increased macromolecular crowding (Extended data Fig. 2h). We also found that droplets diffused more slowly after osmotic compression compared to control (Fig. 2e). Droplet diffusivity depends upon the environment and droplet size. However, average droplet total intensities were lower in osmotic compression conditions, indicating that the reduction in droplet diffusivities was not due to increased droplet size (Fig. 2f). These results support our hypothesis that macromolecular crowding reduces droplet diffusivities and thus inhibits droplet growth.

We next wished to confirm that these effects of the cellular environment were conserved in human cells. To achieve this, we transfected plasmids encoding the two synDrop components into HeLa cells, and observed the formation of droplets (Fig 2g). The amount of DNA delivered to cells is highly variable in transient transfection and, as a result, protein expression levels were highly variable in these experiments. Since protein concentration affects the kinetics of droplet formation, this heterogeneity in protein expression levels across the population made it challenging to study averaged droplet kinetics effectively. Instead, we took advantage of this heterogeneity to define a phase diagram based on the total intensities of two protein components at a fixed time point (Fig 2h).

First, we tested the effects of increasing molecular crowding through osmotic compression. Similar to yeast, we observed a decrease in droplet size and an increase in droplets number per cell after osmotic compression (Fig. 2g, h). We found the synDrops formed in the same region of the phase diagram regardless of experimental condition, suggesting that a specific amount and ratio of protein components is required for synDrop formation, as predicted for a process driven by phase separation (Fig. 2h). However, droplet intensities were lower after osmotic compression compared to control (Fig. 2h, i). Therefore, we conclude that macromolecular crowding inhibits droplet growth in both human and yeast cells.

Next, we employed our agent-based (MD) models to simulate synDrops. These models allowed us to investigate molecular details that are not easily accessible from experimental data. The well-defined structures and binding interactions between the two synDrop components enabled us to quantify droplet network structures with graph-theory based analyses. Here, we defined each synDrop component as a node and the bond between two components as an edge. We calculated the topological shortest distances between each pair of components and mapped out bond connectivity to define each molecular cluster. The distance matrix from this analysis was then used for hierarchical clustering. Within the resulting clustergram, squares along the diagonal correspond to clusters of interacting molecules. Each pixel on the x and y axes represents an interaction between two individual molecules in the simulation system and is colored according to the topological distance between them (e.g., molecules that are directly connected are dark blue, while molecules that are indirectly connected through a chain of interactions are a lighter hue). Blank pixels indicate that there is no path connecting the two corresponding molecules (Fig. 2j and Extended data Fig. 3a).

When there were no crowders in the system, there was very limited cluster formation (Fig. 2j, k – top; Extended data Fig. 3a, b – top; Supplementary Movie). In contrast, large clusters formed when a 30 or 35% volume fraction of crowder was added to mimic the excluded volume typically present in the cytoplasm, suggesting that macromolecular crowding is crucial to nucleate and stabilize synDrop mesoscale networks (Fig. 2j – middle; Extended data Fig. 3a – middle; Supplementary Movie). However, when we further increased the crowder volume fraction to 50% (mimicking crowder concentrations in osmotically compressed cells), we observed a larger number of smaller droplets (Fig. 2j – bottom; Extended data Fig. 3a – bottom; Supplementary Movie). Similar results were also obtained by tracking the number of molecules within the largest cluster (Fig. 2k and Extended data Fig. 3b) and by plotting the cluster size distribution at the end time point (Extended data Fig. 3d). The initial growth rate of average cluster size increased with crowder volume fraction (Fig. 2l and Extended data Fig. 3c), suggesting that nucleation was promoted by macromolecular crowding. However, under high crowding conditions (e.g., 50% volume fraction) cluster size was limited at late time points. These results are consistent with our experimental data that physiological crowding (∼30-35%) appears to be optimal for synDrop assembly. Molecular crowding plays contrasting roles in droplet nucleation and growth. While it is crucial for droplet nucleation, it also inhibits droplet growth.

We next wondered if MD simulations could provide further insights into the molecular basis of the frustration of synDrop growth when there is excessive macromolecular crowding. We plotted the average diffusivity for each cluster as a function of the cluster size and found that diffusivities decrease as a function of cluster size as expected, and were reduced overall when crowder volume fractions were increased (Fig. 2m and Extended data Fig. 3e), consistent with our hypothesis that crowding frustrates coalescence by reducing cluster diffusivities. This effect is particularly pronounced under conditions of excess macromolecular crowding.

Finally, we investigated the molecular basis of the promotion of droplet nucleation by macromolecular crowding. We hypothesized that increased macromolecular crowding could favor binding interactions, as previously reported^8^. To assess this idea, we performed MD simulations on a simplified system where two protein components each had only a single available binding site (1,1) (Extended data Fig. 3f). We then extracted the effective dissociation rate (K_d_) under different volume fractions of crowders by quantifying the number of bonds formation at equilibrium. The effective K_d_ was indeed reduced (affinity was increased) in simulations with increased crowder volume fractions (Fig. 2n and Extended data Fig. 3g). Furthermore, we determined the unbinding rate (k_off_) by allowing the system to reach equilibrium, switching the binding rate to zero, and then measuring the rate at which bonds dissociate (Extended data Fig. 3f). Calculation of effective K_d_ and k_off_ allowed us to infer the effective binding rate (k_on_), which is k_off_ divided by K_d_. Interestingly, when comparing conditions with 30% volume fraction crowder to those with no crowders, we found k_off_ did not change (Extended data Fig. 3f). This indicates that the decrease in effective K_d_ is mainly attributed to an increase in effective k_on_, potentially facilitated by the increase in effective concentration.

In conclusion, our combination of *in vivo* experiments and simulations support the model that macromolecular crowding promotes droplet nucleation by reducing the effective K_d_ of chemical bond formation, but also inhibits droplet growth by reducing droplet diffusivity, therefore kinetically frustrating coalescence of small droplets into larger structures.

## ATP-dependent cellular activities promote droplet growth

We next wondered if other features of the cytoplasmic environment could impact synDrop assembly. In addition to being crowded, the cytoplasm is also far from equilibrium due to ATP-dependent activities. Cellular metabolism was previously shown to strongly affect the motion of mesoscale particles^37^. We therefore hypothesized that ATP-dependent cellular activities might affect synDrop formation by promoting their motion and therefore driving coalescence of small droplets into larger structures.

We used metabolic inhibitors: 2-deoxyglucose (2-DG) and antimycin to deplete intracellular ATP in yeast cells, taking care to maintain neutral pH within cells and isotonic conditions to avoid osmotic perturbations to cell volume^38^. We depleted ATP at different time points to assess the importance of ATP during different phases of synDrop assembly. We observed that synDrop growth was inhibited within 10 min of ATP depletion (Fig. 3a, top), and synDrop diffusivity was also reduced immediately after ATP depletion (Fig. 3a, bottom). These effects were most apparent when ATP was depleted early during synDrop assembly (Fig. 3b). Therefore, cellular active matter is crucial for both synDrop diffusivity and growth in yeast cells.

**Fig. 3.**
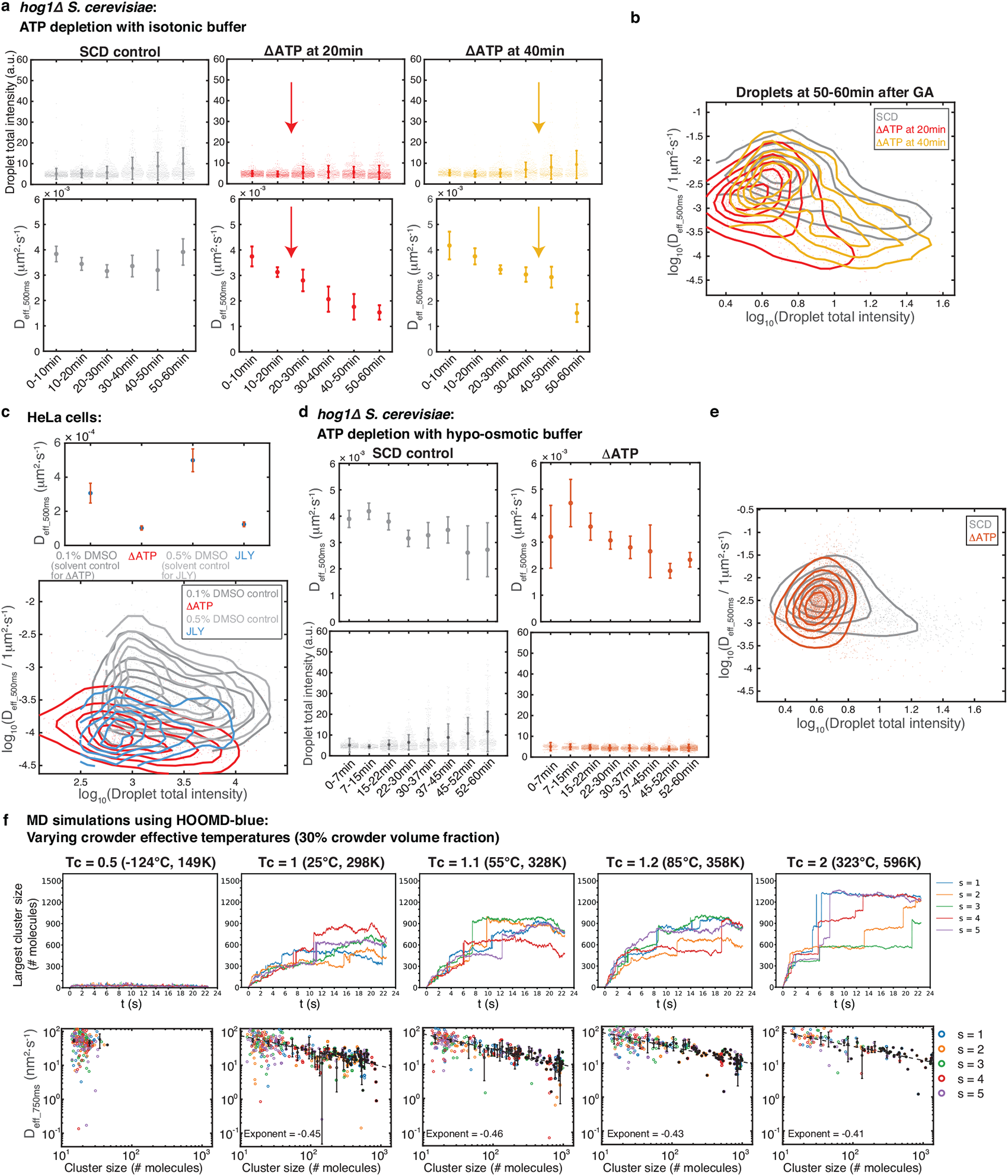
ATP depletion inhibits synDrop growth. **a**, Droplet formation in *hog1*Δ *S. cerevisiae* yeast cells in control conditions and after ATP depletion with isotonic buffer using 80mM sorbitol. ATP was depleted at two time points (indicated by arrows), 20 min and 40 min after synDrop induction with GA: (Top) Droplet total intensity, mean ± SD; (bottom) Median droplet diffusivity ± SEM. **b,** Density plot of individual droplet diffusivity versus its total intensity 50-60 min after induction. **c,** Properties of droplets that were pre-formed by one hour of GA induction with DMSO (solvent control) in mammalian HeLa cells and subsequently treated with the following conditions for one hour: ATP depletion, or the JLY cocktail (which freezes actomyosin dynamics): (Top) Median droplet diffusivity ± SEM; (bottom) Density plot of diffusivity versus total intensity for individual droplets. **d,** Droplet formation in *hog1*Δ *S. cerevisiae* yeast cells comparing control conditions to ATP depletion with a hypo-osmotic buffer: (Top) Median droplet diffusivity. Error bars are SEM; (bottom) Droplet total intensity: averaged values ± SD. **e,** Density plot of diffusivity versus total intensity for individual droplets at all time points post-induction. **f,** MD simulations using HOOMD-blue keeping a constant 30% crowder volume fraction but varying crowder effective temperatures. Values are shown relative to room temperature in units of Kelvin: 0.5, 1, 1.1, 1.2, 2: (Top) Number of molecules within the largest synDrop cluster over time from five replicate simulations; (bottom) Cluster diffusivity versus cluster size (number of molecules). The black data points are the mean of five replicate simulations, error bars are SD, dashed black line is the linear fit in log space with exponent (slope) labeled.

We repeated this experiment in HeLa cells, and found that droplet diffusivity was greatly reduced after we removed all metabolic activity by ATP depletion (Fig. 3c). The dynamics of the actomyosin cytoskeleton are an important source of cellular motion^39^. We therefore hypothesized that actomyosin contractility might agitate the cytoplasm and increase synDrop motion. To test this idea, we inhibited actomyosin dynamics using the JLY drug cocktail, which simultaneously prevents actin depolymerization, polymerization, and myosin II-based restructuring^40^. This treatment reduced diffusivity almost as much as total ATP-depletion, suggesting that actomyosin dynamics is a dominant factor that increases mesoscale diffusivity in the cytoplasm of mammalian cells (Fig. 3c). Depletion of ATP or freezing of actomyosin dynamics using JLY decreased both droplet diffusivity and droplet total intensity (Fig. 3c, bottom); however, there was no clear relationship between droplet local diffusivity and droplet size. In conclusion, actomyosin activity is the dominant ATP-dependent activity that increases synDrop motion in mammalian cells and is required for the formation of large synDrops.

We next wondered if reduced diffusivity of synDrops is the main cause of growth inhibition upon ATP depletion. To test this, we used hypo-osmotic shock to drive water influx and reduce crowding until the diffusivity of synDrops in ATP-depleted cells was the same as that of untreated cells (Fig. 3d above and Fig 3e). However, droplet growth was still inhibited in ATP-depleted cells, even when diffusivity was restored (Fig. 3d, bottom, and Fig. 3e). This result suggested that increasing droplet local diffusivity was insufficient to rescue synDrop growth. Therefore, additional ATP-dependent cellular activities are necessary to promote synDrop growth.

Next, we attempted to model the role of cellular active matter using our MD simulations. To achieve this, we used a simple approximation of altered environmental motion by adding frequency-independent isotropic noise to crowders by varying the effective temperatures of the crowders^41,42^, while keeping the temperature of the synDrop components constant. We observed a positive correlation between the largest cluster size and the effective temperature of the crowders (Fig. 3f, top). When we plotted cluster diffusivity versus cluster size on a log-log scale, we observed individual cluster diffusivities were slightly higher at higher crowder effective temperature conditions and the rate of decrease in cluster diffusivity with increasing cluster size was also slightly slower (Fig. 3f, bottom). However, this increase in cluster diffusivity was relatively modest, implying that other factors may contribute more significantly to the increased mesoscale assembly at higher crowder effective temperature, supporting the experimental results within cells.

## ATP-dependent cellular activities facilitate droplet growth by promoting long-range cellular structural reorganization

Given that coalescence dominates synDrop growth, the growth process is intrinsically linked to droplet motion. Multiple intracellular factors can influence droplet motion, including macromolecular crowding, viscoelasticity, and poroelasticity^43^. Non-equilibrium ATP-dependent cellular activities can modify all of these factors. At small length-scales (< 100 nm), ATP-dependent cellular activities may change the spatial distribution and dynamics of macromolecular crowders. At larger length-scales (> 100 nm), larger cellular structures including membranes and the actomyosin cytoskeleton, both of which undergo dynamic ATP-dependent fluctuations, have strong impacts on mesoscale confinement and elasticity^39,44^. We therefore examined droplet trajectories more closely to gain insight into how longer-range confinement and elasticity relate to synDrop growth.

The average distributions of angles between two vectors that connect subsequent steps in particle tracks can indicate whether particles are driven by active motion, or confined. The angle correlation function is calculated from the ensemble- and time-averaged cosine values of droplet trajectories angles at various time lags^45^. If the average angle between steps is in the range of (π/2 to π) the angular correlation function will be less than zero (-1 to 0). This indicates anti-persistent motion in particle trajectories, suggesting confined or caged particle movement. Conversely, if the averaged angle between steps falls in the range of (0 to π/2), the angular correlation function will be larger than zero (0 to 1). This implies persistent motion in particle trajectories, suggesting motion driven by active processes. We analyzed droplet trajectories at three time points after synDrop induction in control conditions or after ATP depletion in an isotonic buffer in yeast cells (Fig. 4a). In general, we observed negative angle correlations at all time scales, indicating confined motion. This confinement was most apparent at short time-scales. These angle correlation values were further reduced when ATP was depleted, suggesting particles experienced higher confinement. The effect was particularly dramatic when ATP was depleted at later times after synDrop induction when droplets were larger. We conclude that ATP-dependent activities reduce confinement in the cytoplasm and this effect is especially important for larger particles.

We next selected droplets present at 50-60 min after synDrop induction that had similar total intensities. Analyzing this subset of droplets enabled more precise comparison between conditions, and to avoid the concern that the distribution of synDrop sizes changes in different conditions (Extended data Fig. 4a). This approach gave similar results in our angular correlation analyses; ATP depletion led to a reduction in the values of the angular correlation function at all time scales, with more prolonged ATP depletion leading to larger reductions (Extended data Fig. 4b). Therefore, increased confinement after ATP depletion is not a consequence of differences in synDrop size.

**Fig. 4.**
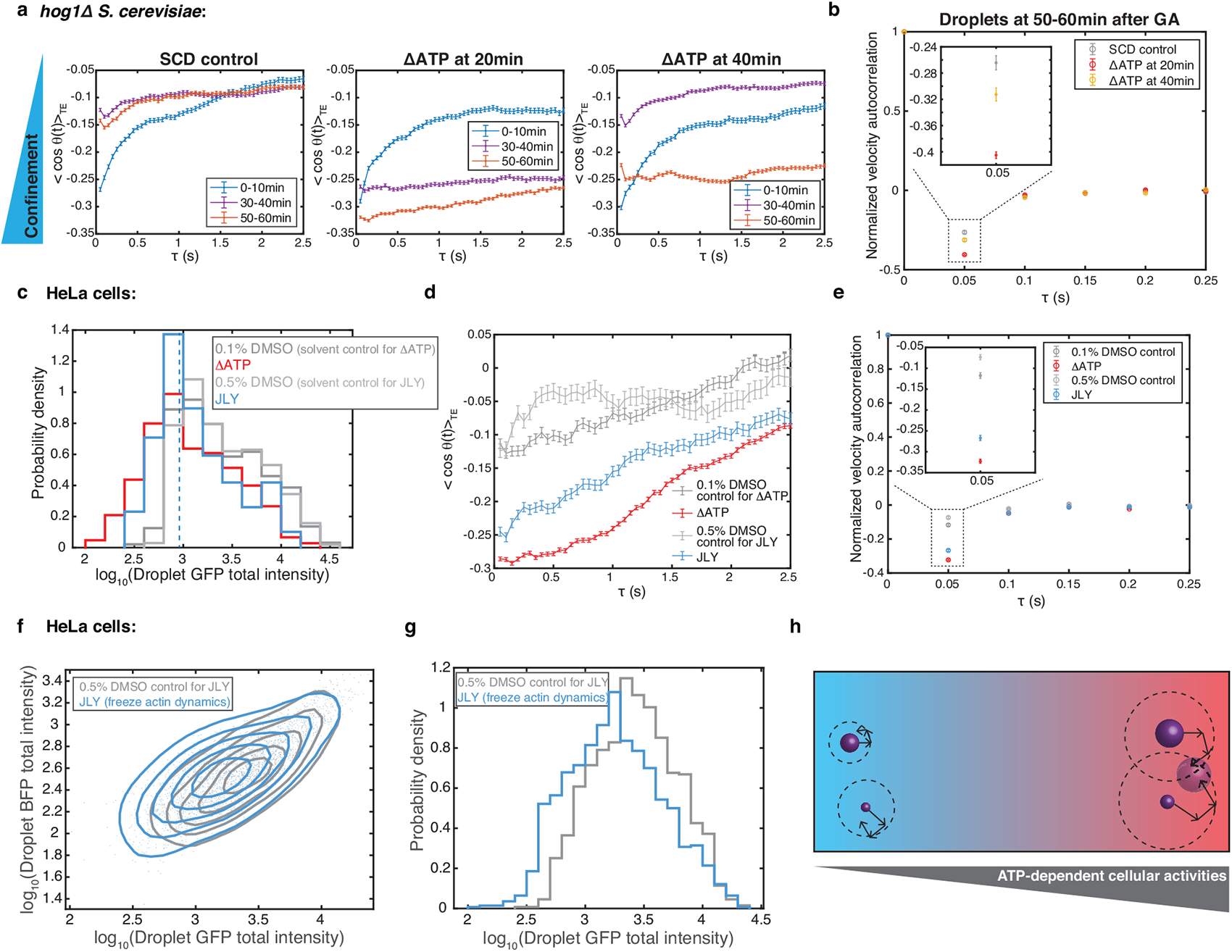
ATP-dependent cellular activity facilitates droplet growth by promoting long-range cellular structural reorganization. **a**, Angle correlation analyses on droplet trajectories at 0-10 min, 30-40 min and 50-60 min after synDrop induction with GA in *hog1*Δ *S. cerevisiae* yeast cells comparing control conditions to ATP depletion at 20 min and 40 min. Lower angle correlation values indicate greater confinement. **b,** Normalized velocity autocorrelation for droplet trajectories in *hog1*Δ *S. cerevisiae* yeast cells. Droplets were analyzed 50-60 min after synDrop induction. **c,** Histogram of droplet GFP total intensity for droplets that had formed after one hour of induction in mammalian HeLa cells comparing control (DMSO) to 1-hour ATP depletion or 1-hour JLY treatment (to freeze actin dynamics) conditions. Droplets with total intensities≤ 900 (blue dashed line) were used in subsequent analyses. **d,** Angle correlation analyses droplets that had formed after one hour of induction in mammalian HeLa cells. Droplets with total intensity ≤ 900 were analyzed after one hour of SynDrop induction. **e,** Normalized velocity autocorrelation for droplets that had formed after one hour of induction in mammalian HeLa cells. **f,** Phase diagram of synDrop BFP and GFP total intensities. Droplets were induced for 1-2 hours in the presence of control (DMSO) or JLY treatment. **g,** Histogram of droplet GFP total intensities (log scale). **h**, Model for how ATP-dependent cellular activities may influence droplet growth.

Finally, we used hypo-osmotic shock to investigate whether equalizing local diffusivity between control and ATP-depleted conditions would also equalize confinement. We found similar patterns of confinement after hypo-osmotic shock; although local diffusivity was equalized, ATP depletion still increased confinement (Extended data Fig. 4c).

Together, these results are consistent with a model where diffusivity of droplets is influenced not only by macromolecular crowding^5^ but also by additional factors that define a longer-range confinement.

We next investigated cytoplasmic elasticity using the velocity autocorrelation function^46^ of the droplet trajectories. We focused on larger droplets that formed 50-60 minutes after synDrop induction. We compared control conditions to ATP-depletion, both in an isotonic buffer (Fig. 4b) and in the hypo-osmotic buffer that normalized local diffusivity (Extended data Fig. 4d). We observed a negative peak, corresponding to anti-persistent particle motion, which is a signature of elastic materials. The magnitude of this negative peak increased after ATP depletion, and this elasticity further increased after longer periods of ATP depletion (Fig. 4b, inset). This suggests that ATP depletion increases the elasticity of the cytosol. This elasticity may impede mesoscale droplet motion and thus impose a longer-range confinement. We conclude that ATP-dependent cellular activities help reduce cellular elasticity, fluidize the cytosol and constantly remodel the cytosol to reduce confinement, leading to increased mesoscale droplet motion, which promotes droplet growth.

We next examined droplet trajectories in mammalian HeLa cells. We focused on a subset of synDrops of smaller size to avoid concerns about differences in droplet size between conditions (Fig. 4c and Extended data Fig. 4e). Similar to the results in yeast cells, angular correlation values were reduced compared to control after ATP depletion (Fig. 4d, red line). Therefore, ATP-dependent activities are required to reduce confinement in mammalian cells.

Next, we inhibited actomyosin dynamics with JLY (Fig. 4d, blue line). Again, we found that droplets were more confined at all time-scales (Fig. 4d). We conclude that actomyosin cytoskeleton dynamics is the dominant ATP-dependent activity that reduces confinement in mammalian cells.

In addition to driving ATP-dependent motion, the cytoskeleton also plays a critical role in determining cellular elasticity^10^. Therefore, we also analyzed velocity autocorrelation before and after ATP depletion or JLY treatment. We found that both treatments led to an increase in the magnitude of the negative peak corresponding to anti-persistent particle motion. Thus, either loss of ATP or inhibition of actomyosin dynamics increases the elasticity of the mammalian cytoplasm (Fig. 4e).

Finally, we examined the effect of JLY treatment on droplet formation in HeLa cells. We compared the droplet phase diagram of control cells to that of cells treated with the JLY cocktail at the same time as synDrop induction. We found that droplet intensities were smaller in JLY-treated cells compared to the DMSO control (Fig. 4f and 4g). Overall, these results support a model in which the actin cytoskeleton promotes long-range structural rearrangements and thereby reduces elasticity and confinement in the cytoplasm. The consequent increase in synDrop motion would promote droplet growth through coalescence (Fig. 4h).

## Discussion

Membraneless organelles carry out many essential cellular functions within cells^1^. Therefore, it is important to understand the spatial and temporal information associated with membraneless organelles formation and dissolution – how cells regulate and coordinate this information. Many studies have focused on specific chemical signals^47^; however, few studies have looked at physical cues, which are indispensable but often neglected. Here, we demonstrated that intracellular macromolecular crowding promotes droplet nucleation by reducing effective dissociation constants of binding reactions but inhibits droplet growth by reducing droplet diffusivities, while ATP-dependent cellular activity promotes droplet growth by fluidizing cellular environment through promoting long-range structural rearrangements.

Macromolecular crowding has several effects on molecular assembly. First, it increases the local concentrations of molecules due to excluded volume occupied by macromolecular crowders^7^. Second, it imposes depletion-attraction forces that increase the propensity of molecular assembly^6^. The cytoplasmic excluded volume is dominated by mesoscale particles, in particular ribosomes, therefore this entropic effect is most prominent at the mesoscale. Both effects can affect binding interactions, leading to reduced effective dissociation constants^8^. Crowding agents have been shown to lower the critical concentrations for several *in vitro* reconstituted phase separation systems^48,49^. However, the inhibition of the kinetics of droplet growth by excess macromolecular crowding is less studied due to the limited availability of controlled *in vivo* phase separation systems.

synDrops have a unique combination of features that make them an ideal platform to investigate how intracellular biophysical environments affect condensates assembly. Droplet nucleation and growth dynamics can be studied on a reasonable time scale (minutes – 1 hour). In contrast to endogenous condensates, the synDrops components were designed to minimize non-specific interactions with endogenous molecules within cells, including ATP-consuming enzymes. Moreover, the well-defined protein structures and network geometry make synDrops highly amenable to simulation and analysis with graph-theoretical approaches.

Our study highlights how the intracellular environment modulates mesoscale molecular assembly through a combination of macromolecular crowding and cellular active-matter. Notably, the intracellular environment is highly heterogeneous in mesoscale diffusivity^50^, reflecting local heterogeneity in macromolecular crowding and cellular activity. These physical variations may underlie the distinct behavior of droplet formation within cells compared to the theoretical prediction that droplets should thermodynamically fuse into a single entity. By actively modulating local macromolecular crowding and cellular activity levels, cells could potentially control the formation of endogenous condensates at different locations via biophysical signals. For example, increased cellular activity, such as actin dynamics near the cell cortex, could facilitate endogenous condensate formation, which might in turn contribute to the nucleation and growth of the cytoskeleton network.

We speculate that changes in the biophysical properties of cells could be sensed by their impacts on condensate assembly. Indeed, a synthetic droplet can modulate the rates of kinase reaction in response to changes in macromolecular crowding, demonstrating the feasibility of this idea in cells^51^. On the other hand, the biophysical properties of the cell interior may also change during disease progression, leading to aberrant phase separation of endogenous condensates. Our study provides a framework to guide future investigations into the effects of intracellular biophysical properties on endogenous condensate formation and dissolution and their relevance to normal biology and disease pathology.

## Acknowledgements

LJH and TS were funded by NIH R01 GM132447, R37 CA240765, the American Cancer Society Cornelia T Bailey Research Award, the NIH Director’s Transformative Research Award TR01 NS127186, the Air Force Office of Scientific Research (AFoSR FA9550-21-1-3503 0091), and the Human Frontier Science Program (RGP0016/2022-102). GMH and GM were supported by the National Institutes of Health (NIH) via Award No. R35-GM138312. This work was supported in part through the NYU IT High Performance Computing resources, services, and staff expertise, and simulations were partially executed on resources supported by the Simons Center for Computational Physical Chemistry at NYU (SF Grant No 839534).

## Methods

### Plasmid construction

For yeast plasmids, open reading frames (ORF) encoding full length Gid and Gai 1-92^26^, dimer (PDB 4LTB) and hexamer (PDB 3BEY)^18^, Green fluorescent protein (Superfolder GFP) and blue fluorescent protein (mTagBFP) were first amplified using PCR. Using Gibson assembly, we then fused each ORF: Gai-BFP-3BEY and Gid-4LTB-GFP with a strong yeast promoter TDH3, an N-terminal nuclear export signaling sequence, and a yeast terminator CYC1, and subsequently assembled into yeast backbone vectors pRS304 and pRS306 respectively. For mammalian plasmids, we codon optimized yeast ORFs based on codon usage of Homo sapiens (Twist bioscience, CA). We then introduced P2A sequence between two ORFs through oligo overhangs and combined Gai-BEY-3BEY and Gid-4LTB-GFP using Gibson assembly onto the same mammalian lentiviral pLVX plasmid with a CMV promoter.

### Yeast transformation

Two yeast synDrops plasmids were first linearized using restriction enzyme cutting within auxotrophic marker region. The linearized plasmids were then transformed into W303 yeast strains (*MATa leu2-3, 112 trip1-1 can1-100 ura3-1 ade2-1 his3-11-,15*) sequentially using a LiAc based protocol^52^. Single yeast cell colony was selected based on whether condensates were able to form after one hour of 300 μM GA induction.

### Mammalian cell transient transfection

Mammalian HeLa cells were plated on a six-well plate in high glucose Dulbecco’s Modified Eagle Medium (DMEM) with L-glutamine (Gibco) supplemented with 10% fetal bovine serum (FBS; Gemini Bio), penicillin (100 U/ml) and streptomycin (100 g/ml) (Gibco). Cells were incubated at 37 with 5% CO_2_ in a humidified incubator and grown to approximately 60-80% confluency after one day of plating. On the next day, cells were transiently transfected using 1g of plasmid and 3ul of FuGENE HD transfection reagent (Promega) based on manufacturer’s protocol. After 24 hours, cells were ready for imaging by replacing with fresh supplemented DMEM. Induction of synDrops in HeLa cells were performed by adding GA till 100 M final concentration.

### Drug treatment

To deplete ATP, S. cerevisiae cells were treated with 20 mM 2-deoxyglucose (2-DG) and 10 μM antimycin A in Synthetic Complete (SC) media without glucose^53^. The media was further supplemented with either 80mM sorbitol for isotonic buffer condition or 10mM sorbitol for hypo-osmotic buffer condition. Additionally, the pH was balanced to 7.5 using a 50mM Tris-HCl buffer. For ATP depletion in mammalian HeLa cells, a mixture of 6 mM 2-deoxyglucose (2-DG) and 1 μM carbonyl cyanide-trifluoromethoxy phenylhydrazone (FCCP) in CO_2_-independent medium supplemented with L-glutamine were added to cells for 1 hour^54^.

To inhibit TOC1 signaling, S. cerevisiae cells were treated with 1 μM rapamycin for 2 hours in Synthetic Complete Dextrose (SCD) media^5^. For JLY treatment, mammalian HeLa cells were first treated with 10 μM y27632 for 10 min, followed by the addition of jasplakinolide and latrunculin B to final concentrations of 10 μM y27532, 8 μM jasplakinolide, and 5 μM latrunculin B^40^ for 1 hour in CO_2_-independent medium supplemented with L-glutamine.

### Microscope imaging of yeast cells

Yeast cells were imaged using TIRF Nikon TI Eclipse microscope with a 100x oil objective (100x phase, NA = 1.4, part number = MRD31901) and a sCMOS camera (Zyla, Andor, part number = ZYLA-4.2p-CL10). Epifluorescence LED light source (Spectra X, part number = 77074160) was used for imaging yeast cells with synDrops. The GFP channel was imaged through GFP filter set (EF-EGFP (FITC/Cy2), Chroma, part number = 49002), while the BFP channel was imaged through quad-band filter set (ET – 405/488/561/640 nm, Chroma, part number = TRF89901). Z stacks were taken for each channel with an interval of 0.5 *μm* and total distance of 3 *μm* (7 slices). Average projection of Z stacks was used for subsequent imaging analyses. Droplet movies were also recorded using GFP channel with 50 ms frame rate without delay for a total of 20 s (total 400 frames). 100% power of 488 nm laser light source (OBIS 100mW LX 488 nm, Coherent, part number = 1236444) with GFP filter set (EF-EGFP (FITC/Cy2), Chroma, part number = 49002) was used for imaging yeast cells with 40 nm diameter Genetically Encoded Multimeric nanoparticles (GEMs). To record GEM movement, we used Highly inclined thin illumination (HILO) mode. Each GEM movie was imaged on a single focal plane at 10 ms frame rate with no delay (100 Hz) for a total of 4 s (total 400 frames).

### Microscope imaging of mammalian cells

Mammalian cells with synDrops were imaged using confocal Nikon TI Eclipse microscope with a spinning disk unit (CSU-X1 spinning disk, Yokogawa, part number = 99459), under a 60x oil objective (60x, NA = 1.49, part number = MRD01691) and a sCMOS camera (Prime 95B, Teledyne Photometrics). The GFP channel was excited through laser light source (OBIS 100mW LX 488 nm, Coherent, part number = 1236444) and imaged through GFP emission filter (EF525/36m, Chroma, part number = 77014803). The BFP channel was excited through LED light source (X-Cite 120LED, Excelitas, part number = 010-00326R) and imaged through DAPI filter set (ET-DAPI, Chroma, part number = 49028). Z stacks were taken for each channel with an interval of 0.5 *μm* and total distance of 6 *μm* (13 slices). Average projection of Z stacks was used for subsequent imaging analyses. Droplet movies were also recorded using GFP channel with 50 ms frame rate without delay for a total of 20 s (total 400 frames).

### Image analyses

To characterize synDrops’ properties within cells, images after average Z projections were analyzed using the Trackmate plugin^55^ on ImageJ^56,57^. Due to higher signal-to-noise ratio in GFP channel compared to BFP channel as well as droplets overlap in GFP and BFP channels, droplets were then only detected using GFP channel. We applied a LoG (Laplacian of Gaussian filter) detector on Trackmate to identify the droplets, with one micron ‘Estimated object diameter’ and a fixed ‘Quality threshold’ across all different conditions in each experiment. To determine the droplet total intensity, we employed a circular particle detection algorithm that identified each droplet with a fixed circular region larger than the droplet itself. Subsequently, we computed the mean pixel intensity within this identified region and subtracted the background mean pixel intensity. We denoted this measurement as the droplet total intensity because it reflects both the number of molecules and molecular concentration within the condensate. Results of particle detection from Trackmate included all time points and were then saved as *xlm* files. Using home-written MATLAB (2019a) code, we subsequently extracted and compiled droplets information, including droplet raw mean pixel intensities and their locations in the images. We also determined the background mean pixel intensity by randomly selecting 20 circles with the same one-micron diameter in each image from areas away from droplets. Thus, final droplet total intensities were calculated by subtracting the background mean pixel intensity from droplet raw mean pixel intensities identified above. Yeast-Spotter^58^ were used to identify single yeast cells, which generated ImageJ mask files indicating locations of each individual yeast cell. By combining with droplets’ location information, we computed the number of droplets per cell using home-written MATLAB code.

To obtain droplet diffusivities, we used simple linear assignment problem (LAP) particle tracking function on Trackmate in addition to particle detection, with max linking and gap-closing distance of 390 nm and maximal gap-closing frame interval of 1. Only trajectories with more than 10 time points were included for subsequent mean squared displacement analyses using home-written MATLAB code.

To determine GEM diffusivities, GEM trajectories were detected using the Mosaic plugin^59^ on ImageJ^56,57^ with particle detection parameters of radius 3, cutoff 0, per/Abs (percentile) 0.1 and particle linking parameters of link range 1, displacement 5 with Brownian dynamics. We only selected trajectories with more than 10 time points for subsequent mean squared displacement analyses using home-written MATLAB code.

To generate density plots based on the scattered data points, the data space was separated into 25 x 25 different regions based on the minimum and maximum values on both x and y axis. By counting the number of data points within each region, 2D density matrices were generated and smoothed using the MATLAB function scattercloud^60^. The contour lines were further obtained using MATLAB function contour.

### MD simulation analyses

We used graph-theory based methods for analyzing MD simulations. Each molecule within MD simulations had a unique number identifier and was treated as the node for the graph. Bonds formed at each time point were recorded based on molecule pairs that formed each bond, and were treated as the edges for the graph. The graph at each time point was then constructed by providing both node and edge information inputs using the igraph^61^ package in python. To identify clusters, a distance matrix was first calculated based on the topological shortest path that links each pair of molecules. Subsequently, a hierarchical clustering algorithm was employed on the distance matrix. This led to the reordering of molecule sequences, with molecules within each cluster being grouped together. Cluster size was then determined based on the number of molecules that were within each cluster. Locations of each molecule were also recorded at each time point.

To calculate cluster diffusivity, clusters with size larger than 10 molecules were first identified at each time point. Pairwise clusters from consecutive time points were connected from the last time point by determining the largest number of same nodes, thus forming trajectories. If a cluster’s size changed by 20 within a time interval, it was considered as a new cluster and tracked as a distinct trajectory. Only trajectories with more than 10 time points were selected for calculating cluster diffusivities, where mean squared displacement of the cluster’s center of mass for each trajectory was fitted over the first 10 time intervals. All analyses were performed using home-written python3 code.

To determine the effective dissociation constants (K_d_) of the chemical bonds, we analyzed the kinetics of bond formation in monomeric MD simulations until equilibrium was achieved. In the monomeric system (where the dimers and hexamers are in a 1:1 stoichiometric ratio), we reduced the available binding sites of two components from 6 and 2 to 1 and 1 each. By fitting the data to an exponential decay function, we extracted the number of bonds formed at equilibrium. Subsequently, K_d_ was calculated based on the concentration of all species at equilibrium, using the formula for a dimerization reaction *A* + *B* ↔ *AB*,
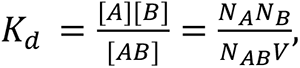 where V is the volume of the simulation box.

To roughly match simulation time scales to experimental ones, the mean-squared displacement of 40 nm GEM particles in simulation were fit to the Einstein diffusion relation in 3d *MSD*(*t*) = 6*Dt* for long times, and *D* was obtained in the units of *μm*^2^/τ. The unit of time τ = **7.5** × **10**^−**8**^ seconds was then obtained by matching this *D* to an approximate cellular value of *μm*^2^/s.

### Simulations set up using HOOMD-blue

An agent-based molecular dynamics (MD) simulation approach has been developed to study the synDrops system. MD Simulations were performed using HOOMD-blue v2.9.6 ^28,29^, making use of a single graphics processing unit (GPU) to achieve considerable acceleration in simulation speeds. We use coarse-grained (CG) representations of each synDrops component—(i) a sphere with six rigid evenly distributed binding sites to represent the hexamer and (ii) 3 spheres in a rod-like arrangement with two complementary binding sites at two ends to represent the rigid coiled-coil dimer. We have 1170 dimers and 390 hexamers within a cubic box with 860 nm sides (maintaining a 3:1 stoichiometric ratio of dimers and hexamers to have 1:1 ratio of complementary binding sites). This results in concentrations of 3 *μM* for dimers and 1 *μM* for hexamers, similar to our estimated values in experiment. Finally, 20 spheres of diameter 40 nm are added to mimic the trace presence of GEMs in the experiment.

In addition, spherical components of various sizes without any binding site are added in the system to mimic the crowded cellular environment. For the initial configuration, the CG components are arranged in a lattice whose positions are generated from a CsCl-type lattice generator using the ‘lattice’ module from the ASE (Atomic Simulation Environment) ^62^ package. We ran MD simulations with varying volume fractions of ribosomes to study the effect of crowding in synDrops assembly. We also varied the effective temperatures that only govern the ribosome movements to study how cellular activities affect synDrops assembly.

Binding occurs through complementary interaction sites between dimers and hexamers. We modeled such covalent interactions by developing an open-source C++ plugin, called the Dynamic Bond Updater ^30^ in HOOMD-blue that builds upon a model for epoxy binding developed in ^63^. The Bond Updater, for every n steps during the MD simulation, stochastically adds or removes dynamic bonds. Binding events occur with a fixed probability P_on_ at a critical distance d_bind_ between interaction sites, while unbinding events occur with a probability P_off_. Using our dynamic bonding framework, we thus have controls over our binding and unbinding rate constants k_on_ and k_off_ respectively; the bond affinity ε is defined by

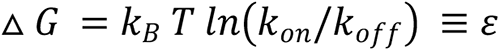

and can be increased by lowering the unbinding rate constant k_off_. We ensure that the dynamic bonding model satisfies detailed balance using a particular Metropolis-like criterion ^64–66^, so that the system moves towards an equilibrium ensemble as bonds form and dissolve dynamically. We use the cell neighbor list ^67^ to accelerate non-bonded agents’ calculations and possible bonding pairs’ constructions.

Interactions between crowders and synDrops proteins occur via a soft repulsion potential ^30^ defined by

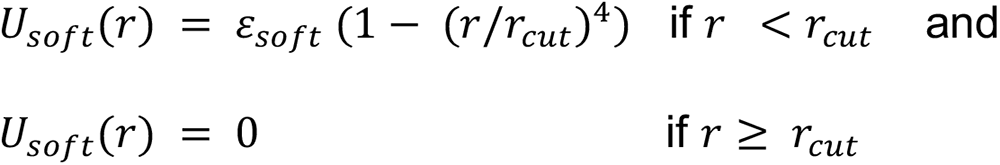

where smoothing was applied using HOOMD-blue’s XPLOR ^29^ smoothing function. The soft potential was implemented by using HOOMD-blue’s tabulated potential option (with 1000 interpolation points between r_min_ = 0 and r_max_ = 1.5σ, where σ is the sum of the radii of the particles). Here, r_on_ is chosen as the point at which the smoothing starts.

We set r_on_ = 0.95r_cut_ for our simulations, and r_cut_ = σ. There is no soft repulsion between complementary binding interaction sites on hexamers and dimers, where we implemented a Lennard-Jones (LJ) ^68–71^ attraction between the hexamer and dimer rigid bodies, with a cut-off distance equal to 2.5σ.

All objects in the system undergo thermal fluctuations using Langevin ^72^ dynamics, with drag forces proportional to the diameter. The dimensions of every CG component approximate their respective crystal structures. Within our MD simulations, we typically use periodic boundary conditions (PBC). However, we also have the option of adding ‘walls’ to confine our system in a ‘closed box’. For volume fractions up to 35%, we are able to place ribosomes in the box without overlaps through random sequential insertion. For higher concentrations, we first set up initial simulation box size using lengths of 1400 nm on a side (4.3x the target volume) and the appropriate number of ribosomes, and then compress the system to the target size of 860 nm linearly over 5 × 10^5^ simulation steps (using the ‘hoomd.variant’ module of HOOMD-blue), and finally turn on dynamic bonding in the system to record synDrops dynamics.

To study how non-thermal cellular activity^41,42^ impacts formation of synDrops via MD simulations, we assign the crowders a different effective temperature T_c_ from the rest of the system, which can be achieved through separate Langevin ‘thermostats’ in HOOMD-blue. We ran a different set of MD simulations at a fixed volume fraction of ribosomes (= 30%) but varying the crowder effective temperatures T_c_.

**Table of parameters for the MD Simulations using HOOMD-blue:**

| Parameter (with description) | Value used in simulations |
| --- | --- |
| Simulation timestep ($dt$) | 0.002 |
| Crowder Temperature $T_c$ relative to $T_{room} = 298.15K$ | 0.5 - 1.0 |
| Simulation box length (in nm) | 860 (for the actual system)<br>400 (for the monomeric system to obtain $K_d$) |
| Number of rod proteins | 1170 (for the actual system)<br>200 (for the monomeric system) |
| Number of hexamers | 390 (for the actual system)<br>200 (for the monomeric system) |
| Number of GEMs | 20 |
| Number of binders on each hexamer | 6 |
| Number of binders on each rod | 2 |
| Volume fraction of ribosomes (crowders) | 0.0 - 0.5 |
| Diameter of inner rod particle (in nm) | 11.7 |
| Diameter of the two outer rod particle(s) (in nm) | 13.45 |
| Diameter of hexamer (in nm) | 12.6 |
| Diameter of ribosome (in nm) | 30.0 |
| Diameter of GEM (in nm) | 40.0 |
| Diameter of binders on rods and hexamers (in nm) | 2.0 |
| Maximum binding distance $d_{bind}$ (in nm) | 1.0 |
| Repulsion for soft quartic potential (in $k_B T$) | 500 |
| $\varepsilon$ for Lennard-Jones potential (in $k_B T$) | 0 |
| Rate constant for dynamic binding $k_{on}$ (in units of $1/\tau$) | 50.0 |
| Rate constants for dynamic unbinding $k_{off}$ (in units of $1/\tau$) | 0.001, 0.0001 |
| Binding affinity $\varepsilon$ (in $k_B T$) | 10.8, 13.1 |

### Simulations set up using custom-developed Java program

A custom 3-dimensional agent-based Java program was developed to simulate aggregation and cluster formations of proteins in a cellular environment. All objects in these simulations are spheres, or spherical aggregates, that move in space as a result of applied forces. These forces arise in three distinct ways: through collisions with other spheres and with the boundaries of the simulation, through bonds to other spheres, and through a random force and torque applied to approximate the Brownian motion of each object. The movements of all molecules then follow Langevin dynamics with a defined effective temperature.

Collisions are resolved with a simple rule: at a low-Reynolds number we can calculate the exact force to resolve any pairwise collision. This method is described in detail in Alberts & Odell (2004)^73^. All the pairwise forces are summed and then attenuated for numerical stability such that collisions resolve over several time-steps (not instantaneously). Translational bond forces are resolved in the same way: by calculating the force required to bring a stretched bond back to its relaxed position. By contrast, torques are calculated with a linear torsional spring. Brownian forces and torques are randomly taken from a Gaussian distribution with a mean of zero and variance of 2DΔt, where D is the diffusivity of the object (which can differ in all three translational and rotational degrees of freedom) and Δt the time-step.

The two protein components of the synDrops system were modeled as spheres having 6 or 2 uniformly distributed binding sites on their surface (Extended data Fig. 1c). The size of each sphere was determined based on its experimental correspondence with known protein crystal structures. Bond kinetics in the model arise by prescribing binding and unbinding rates, with binding occurring between available sites only when they are within a minimum distance of each other. The unbinding rate is assumed to be independent of any strain in the bond. After selecting appropriate values for minimum binding distance and unbinding rate, we ensured that the dissociation constant for the chemical bond ranged between 1 to 10 *μM*. As in the HOOMD-Blue system, we use 1170 dimers, 390 hexamers, and varying numbers of ribosomes in a box with side length 860 nm.

**Table of parameters for the MD Simulations using custom-developed Java program:**

| Parameter (with description) | Value used in simulations |
| --- | --- |
| Simulation timestep ($dt$) (in second) | $10^{-7}$ |
| Crowder Temperature $T_c$ relative to $T_{room} = 298.15K$ | 1 |
| Viscosity (in $Pa \cdot s$) | 0.03 |
| Simulation box length (in nm) | 860 (for the actual system)<br>400 (for the monomeric system to obtain $K_d$) |
| Number of rod proteins | 1170 (for the actual system)<br>200 (for the monomeric system) |
| Number of hexamers | 390 (for the actual system)<br>200 (for the monomeric system) |
| Number of binders on each hexamer | 6 |
| Number of binders on each rod | 2 |
| Volume fraction of ribosomes (crowders) | 0.0 - 0.5 |
| Diameter of rod (in nm) | 23.4 |
| Diameter of hexamer (in nm) | 12.6 |
| Diameter of ribosome (in nm) | 30.0 |
| Maximum binding distance $d_{bind}$ (in nm) | 2.3 |
| Rate constants for dynamic unbinding ($k_{off}$) (in $s^{-1}$) | 26 |

