## Supplemental Figures for "Mesoscale molecular assembly is favored by the active, crowded cytoplasm"

**Extended data Fig. 1: synDrops in mammalian HeLa cells, and alternative agent-based MD simulation platform.**

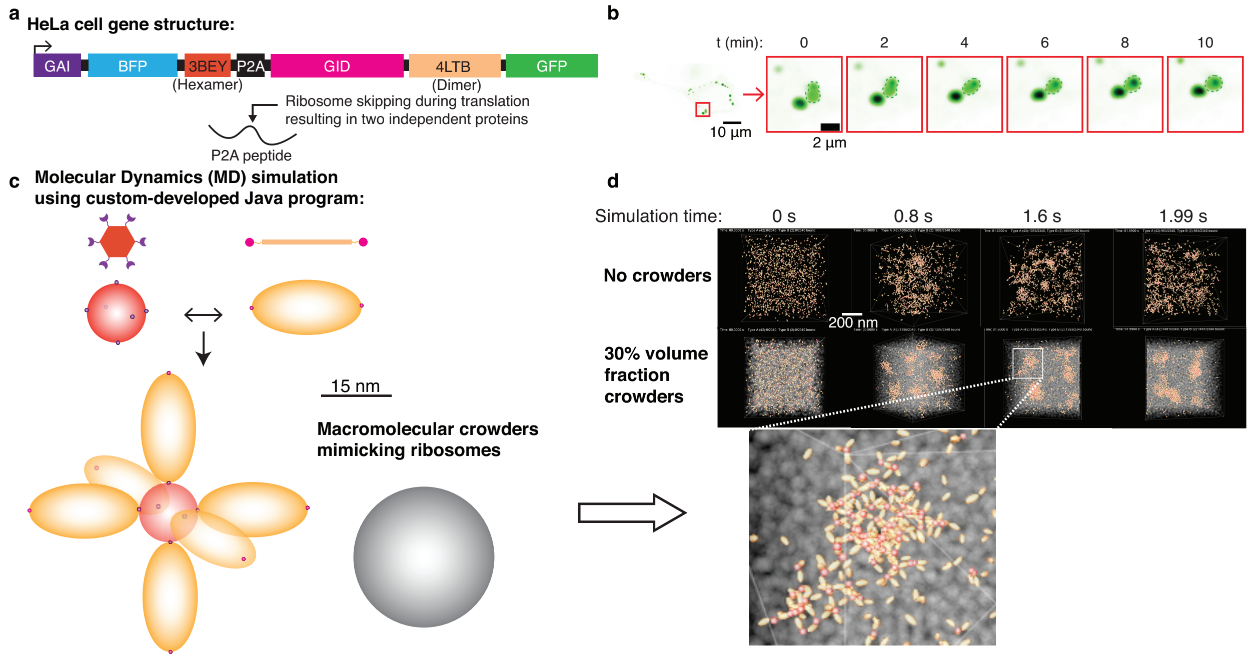


**Extended Data Fig. 1 synDrops in mammalian HeLa cells, and alternative agent-based MD simulation platform. a,** To efficiently express the two protein components of the synDrop system at similar levels in mammalian cells, the two open reading frames (ORFs) were connected by a P2A sequence. The two ORFs are translated one after the other from a single transcript due to ribosome skipping. **b,** synDrops fuse within minutes, suggesting liquid-like properties. **c,** A second MD simulations platform was developed based on a custom Java program. The two protein components of the synDrops system were modeled as spheres with either 6 or 2 binding sites. The simulation system also includes a third molecular component, without binding sites, that mimics ribosomes as macromolecular crowders. **d,** MD simulations of synDrop assembly over time without crowders (top) and with 30% volume fraction of crowders (bottom).

**Extended data Fig. 2: Experimental results: Increasing molecular crowding promotes synDrop nucleation but inhibits mesoscale growth.**

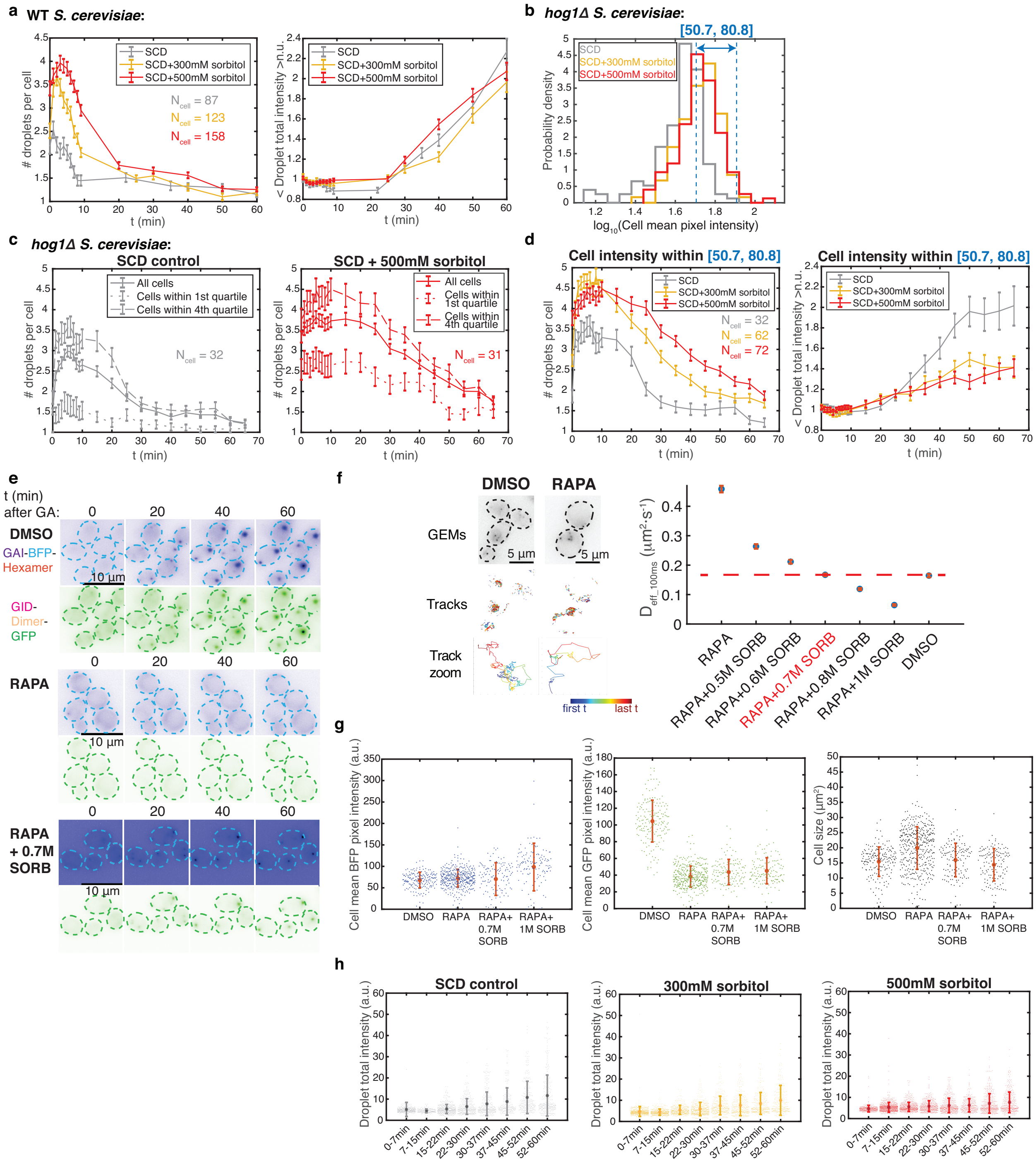

**Extended Data Fig. 2 Experimental results: Increasing molecular crowding promotes synDrop nucleation but inhibits mesoscale growth. a,** Average number of droplets per cell (left) and normalized droplet total intensity (right) during one hour of synDrop induction in WT *S. cerevisiae* yeast cells comparing control to increased molecular crowding conditions (osmotic compression with 300 mM or 500 mM sorbitol). **b,** Osmotic compression increases protein concentrations, but this effect can be accounted for through selection of a subset of cells. Distributions of GFP intensities (mean pixel fluorescence intensities) of *hog1Δ S. cerevisiae* comparing control to osmotic compression conditions. The 4th quartile of GFP intensities in control conditions is labeled by blue dashed lines. **c,** Average number of droplets per *hog1Δ S. cerevisiae* cell after synDrop induction in control (left) and osmotic compression (500 mM sorbitol, right) conditions comparing: all cells, cells with GFP intensities in the lowest ($1^{st}$) quartile, and cells with GFP intensities in the highest ($4^{th}$) quartile. Error bars are SEM. **d,** Kinetics of formation of synDrops in cells with GFP intensities in the $4^{th}$ quartile of control conditions (blue dashed lines in **b**) comparing control to osmotic compression (300 mM or 500 mM sorbitol) conditions. Error bars are SEM. **e,** Representative images of synDrops during one hour after synDrop induction with GA comparing *hog1Δ S. cerevisiae* yeast cells treated with: DMSO (control), rapamycin pre-treatment for 2h (RAPA), and rapamycin pre-treatment for 2h followed by osmotic compression to restore Genetically Encoded Multimeric (GEM) diffusivity to control values (RAPA + 0.7 M Sorb). **f,** GEM nanoparticles were used to determine the sorbitol concentration that restores mesoscale crowding in cells pre-treated with RAPA for 2h to a level comparable to DMSO control cells. **g,** Quantifications of cell mean BFP (left) and GFP (middle) pixel intensities, and cell size (right) comparing *hog1Δ S. cerevisiae* yeast cells treated with: DMSO control; RAPA; RAPA followed by 0.7 M sorbitol; and RAPA followed by 1 M sorbitol. Error bars are SD. **h,** Average droplet total intensity at different time points after synDrop induction under the same conditions as **Fig. 2e**. Error bars are SD.

**Extended data Fig. 3: Simulation results from Java-based MD simulations: Increasing molecular crowding promotes synDrop nucleation but inhibits mesoscale growth.**

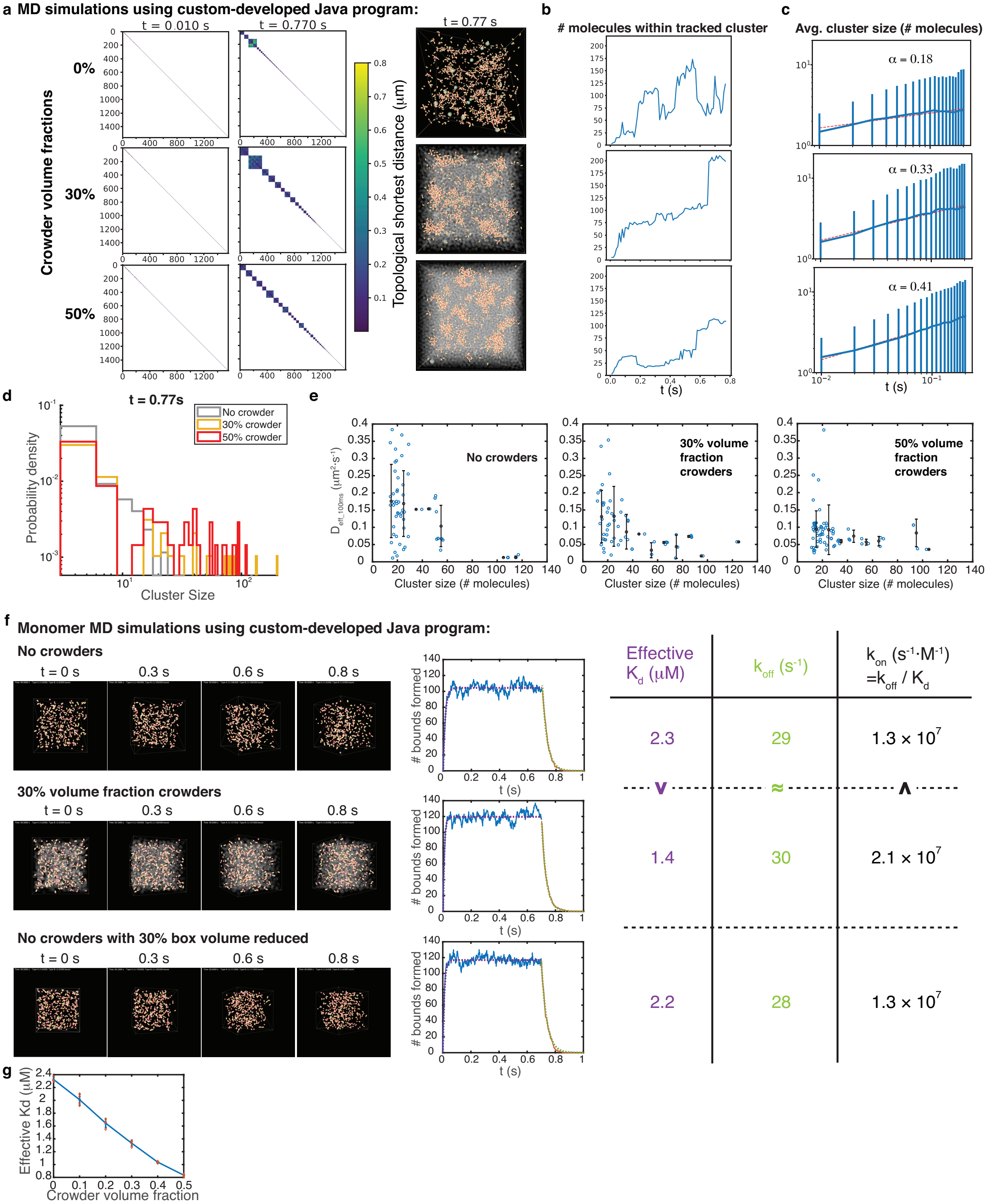


**Extended Data Fig. 3 Simulation results from Java-based MD simulations: Increasing molecular crowding promotes synDrop nucleation but inhibits mesoscale growth. (a-e),** Analyses of MD simulations with 0%, 30% and 50% volume fraction of crowder: **a,** Left) Clustergrams representing molecular connectivity, determined by graph theory. Squares on the diagonal are indicative of clusters of molecules. Analyses are shown after 0.01 s and 0.77 s simulation time. Right) images of the simulations after 0.77 s. **b,** Number of molecules within the largest cluster as a function of time. **c,** Average cluster size (number of molecules) as a function of time. Dashed line is the power law fit with fitted exponent labeled as $\alpha$. Error bars are standard deviation (SD). **d,** Distribution of cluster sizes (number of molecules) at t = 0.77s. **e,** Average cluster diffusivity versus cluster size (number of molecules). Error bars are SD. **f,** Monomer MD simulations to determine binding rates. Each protein component was modified to be monovalent and simulations were performed with no crowders (top), 30% volume fraction of crowders (middle) and no crowders but with a 30% reduction in 30% box volume (bottom). After 0.7s, the binding probability was set to 0 allowing determination of unbinding rates. Effective binding rates ($k_{on}$), unbinding rates ($k_{off}$), and dissociation constants ($K_d$) were inferred from the number of bonds formed over time (table, right). **g,** Effective dissociation constants ($K_d$) as a function of crowder volume fractions. Error bars are SD from three repeats.

**Extended data Fig. 4: ATP-dependent cellular activity facilitates droplet growth by promoting long-range cellular structural reorganization**

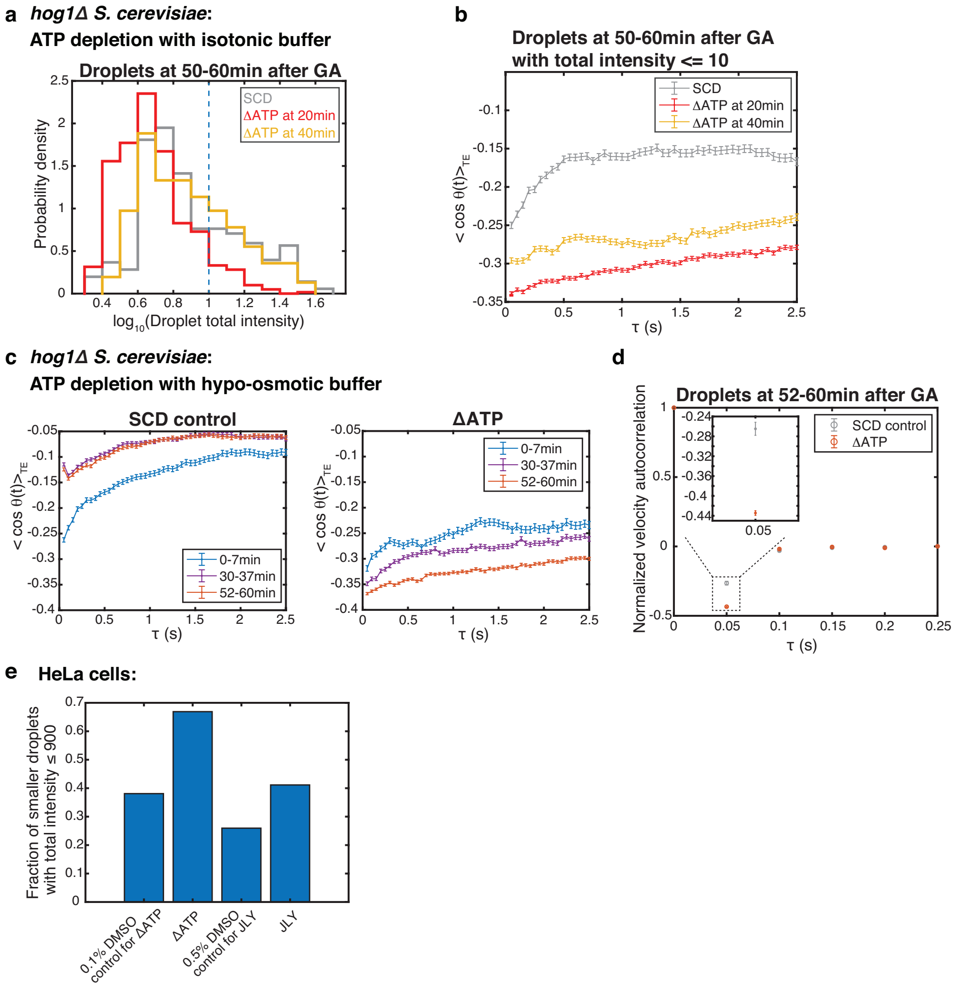


**Extended Data Fig. 4 ATP-dependent cellular activity facilitates droplet growth by promoting long-range cellular structural reorganization. a,** Histogram of droplet GFP total intensity 50-60 min after synDrop induction with GA in *hog1Δ S. cerevisiae* yeast cells in control conditions or after ATP depletion with isotonic buffer at 20 min and 40 min after synDrop induction. Droplets with a total intensity ≤ 10 (blue dashed line) were analyzed in **b**. **b,** Angle correlation analyses of droplets with total intensity ≤ 10. **c,** Angle correlation analyses of droplets trajectories 0-7min, 30-37 min, and 52-60 min after synDrop induction in *hog1Δ S. cerevisiae* yeast cells comparing control to ATP depletion with hypo-osmotic buffer (simultaneous with synDrop induction). **d,** Normalized velocity autocorrelation for droplet trajectories 52-60 min after synDrop induction in *hog1Δ S. cerevisiae* yeast cells using the same conditions as **c**. **e,** Fraction of droplets with total intensity ≤ 900 in various conditions. Droplets were pre-formed by one hour of GA induction with DMSO (solvent control) in mammalian HeLa cells and subsequently treated with the indicated conditions for one hour.

**Supplementary movie Molecular dynamics simulations were performed using HOOMD-blue to compare conditions with 0%, 35% and 50% volume fractions of crowders.** Top) Simulation renderings of cluster formation at all time points. Bottom) Graph theory analyses of cluster formation at the corresponding time points as shown in the top renderings. Clustergrams were employed to illustrate molecular connectivity, where squares along the diagonal represent molecular clusters.
